# Plasticity of behavioural variability reflects conflicting selection in group-living and solitary desert locusts

**DOI:** 10.1101/2020.03.18.996983

**Authors:** Ben Cooper, Jonathan M. Smith, Tom Matheson, Swidbert R. Ott

**Affiliations:** University of Leicester, Department of Neuroscience, Psychology and Behaviour, University Road, Leicester, LE1 7RH, UK

## Abstract

Animals living in groups tend to express less variable behaviour than animals living alone. It is widely assumed that this difference reflects, at least in part, an adaptive response to contrasting selection pressures: group-living should favour the evolution of more uniform behaviour whereas lone-living should favour behaviour that is less predictable. Empirical evidence linking these contrasting selection pressures to intrinsic differences in behavioural variability is, however, largely lacking. The desert locust, Schistocerca gregaria, manifests in two very distinct eco-phenotypes, a lone-living cryptic “solitarious phase” and a swarming “gregarious phase” that aggregates into very large and dense groups. This “phase polyphenism” has evolved in response to contrasting selection pressures that change rapidly and unpredictably. Phase differences in mean behaviour are well-characterised, but no previous study has considered differences in variability. Here we used locust phase polyphenism to test the hypothesis that group living leads to the evolution of reduced intrinsic variability in behaviour. We measured two behaviours in both phenotypes: locomotor activity in the presence of conspecifics, and locomotor hesitation in approaching food when alone. We assayed each individual repeatedly and estimated variability relative to the mean in log-normal mixed-effects models that explicitly account for the means-variance dependency in the behavioural measures. Our results demonstrate that relative behavioural variability differs between the two phases in line with predictions from ecological theory: both within-individual and between-individual variability were lower in the group-living gregarious phenotype. This contrasts with previous studies on social niche construction in spiders and crickets, and highlights the importance of social ecology: in animals that form non-social collectives, such as locusts, reduced individual behavioural variability is key for coherent collective behaviour. The differences in variability persisted when gregarious locusts were tested in isolation and solitarious locusts were tested in groups, indicating that they arise not simply as flexible reactions to different social contexts, but are intrinsic to the individual animals of each phase. This “variance polyphenism” in locusts provides empirical evidence that evolutionary adaptation for group living has driven a reduction in within- and between-individual behavioural variability.

## Introduction

The behaviour of animals varies over time and differs between individuals ^[1]^. Animals living in groups tend to show less behavioural variability than those living alone^[2]^. This reduced variability in groups is thought to be adaptive with respect to predation; group members may benefit from behaving more uniformly as predators disproportionately target individuals whose behaviour differs from the majority ^[3]^. Conversely, animals living alone may benefit from behaving in a less predictable manner to confound predators; a mechanism known as protean behaviour ^[4]^. The contrasting selection pressures of group- and lone-living are therefore expected to lead to adaptive differences in the extent of behavioural variability^[5]^.

Reduction in either within- or between-individual variability of behaviour contributes to greater homogeneity of animal collectives^[6]^. Previous studies have largely focussed on changes in behavioural variability as an instantaneously *reactive* process, when animals behave in one way when alone, but immediately adjust their behaviour to be less variable when in the presence of conspecifics^[2]^. These changes often arise through simple local rules governing movement in response to neighbours ^[7]^ and can be interpreted as ‘contextual plasticity’ ^[5]^. Most research into the effect of predation on behavioural variability uses the framework of contextual plasticity ^[8,9]^, with individuals increasing their within-individual variability in the presence of predator cues. However, a framework that is entirely based on context-driven plasticity of within-individual variability makes it harder to clarify the *evolution* of changes in behavioural variability, because contextual plasticity is thought to shield genotypes from selection^[10]^.

An alternative explanation for how changes in behavioural variability occur is that they are an *intrinsic* process, where prior external conditions induce changes in behavioural variability which then become intrinsic to the individual, persisting beyond the triggering conditions^[11–13]^. This implies that the extent of behavioural variability is regulated by intrinsic mechanisms and forms part of an animal’s phenotype ^[14]^. Indeed, very recent studies report evidence for a genetic component to between-individual differences in within-individual variability^[15]^. While such differences in intrinsic behavioural variability are predicted by theory to be subject to selection, empirical evidence for adaptive changes is largely lacking ^[5]^.

Here, we present robust evidence demonstrating that selective pressures have shaped intrinsic behavioural variability in the desert locust *Schistocerca gregaria*. In this species, the same genotype can manifest in two very distinct eco-phenotypes or phases, a lone-living ‘solitarious’ phase and a ‘gregarious’ phase that forms large, dense groups^[16]^. Solitarious individuals show low levels of activity, have cryptic green or beige colouration and avoid other locusts, whereas gregarious individuals are highly active, are attracted to conspecifics, develop a striking aposematic yellow-and-black colouration, and can aggregate into swarms containing billions of individuals ^[17]^.

This capacity for extreme density-dependant phenotypic plasticity (‘phase polyphenism’) evolved in response to conflicting selection pressures on isolated and swarming individuals that change rapidly and unpredictably ^[18]^. We therefore used phase polyphenism to test the hypothesis that group living leads to reduced intrinsic variability in behaviour. By testing crowd- and isolation-reared animals alone, and with a social stimulus present, we specifically examined whether changes in variability outlast the social conditions which initially trigger them. Our results demonstrate that behavioural variability differs between the two phases in line with predictions from ecological theory ^[5]^. Moreover, these differences arise not simply as flexible reactions to different social contexts, but are intrinsic to the individual animals of each phase. We conclude that this intrinsic individual variation in behaviour provides a substrate for the action of natural selection, rather than shielding the animal from it.

## Methods

### Animals

Desert locusts (*Schistocerca gregaria*, Forskål) were obtained from two breeding colonies maintained at the University of Leicester. The ‘Leicester strain’ colony has been maintained in the laboratory for about 40 years with intermittent introduction of animals from other lab or commercial populations; the ‘Mauritanian strain’ was derived from eggs collected in the field in Mauritania in Spring 2016. All experiments used locusts in their final instar stage. A detailed description of locust rearing conditions is provided in the Supplementary Methods.

In brief, gregarious locusts of both strains were reared at densities of 100–300 locusts per 50 cm L × 50 cm W × 50 cm H cages and fed fresh wheat seedlings and wheat bran *ad libitum*. Solitarious locusts were generated from this gregarious stock following Roessingh *et al*. (1993) ^[19]^. Second-generation solitarious animals were kept individually in separate metal cages (10 cm L × 10 cm W × 20 cm H) from which they could not see other locusts. Cages were supplied with conditioned outside air to provide olfactory isolation. Solitarious cages contained a vial of wheat grass (seedling leaves) and a dish of wheat bran which were replaced twice a week.

Beam-walk experiments used solitarious animals from five different females, and Roessingh arena experiments used solitarious animals from 29 different females. A cohort of solitarious ‘breeders’ was set up monthly; members of this cohort were then bred to produce solitarious animals for experiments.

The parental lineages of the gregarious locusts used in experiments were not recorded. Beam-walk experiments used gregarious animals from five different cohorts, and Roessingh arena experiments used gregarious animals from 12 different cohorts. A new cohort of gregarious hatchlings was set up weekly, and cohorts were euthanised after 2–3 months.

### Beam-walk behavioural assay

Locomotor hesitation of individual locusts was assessed in a beam-walking paradigm, in which the locust exited a holding tube onto a horizontal wooden dowel beam and walked upwind in an airstream that carried fresh wheat grass odour (Fig. 1A). The arena was simplified from a published Y-maze used for testing odour preferences ^[20]^. The wooden beam was suspended above the floor of the arena, a white opaque acrylic box with a transparent acrylic lid. Wheat grass was not visible to the locust, but its odour was drawn through a fine cotton mesh screen towards the animal by a fan which created unidirectional airflow. Six identical arenas were located in a temperature- and humidity-controlled room (36°C, 20–25% RH, >20 air changes per hour) and lit from above.

**Figure 1:**
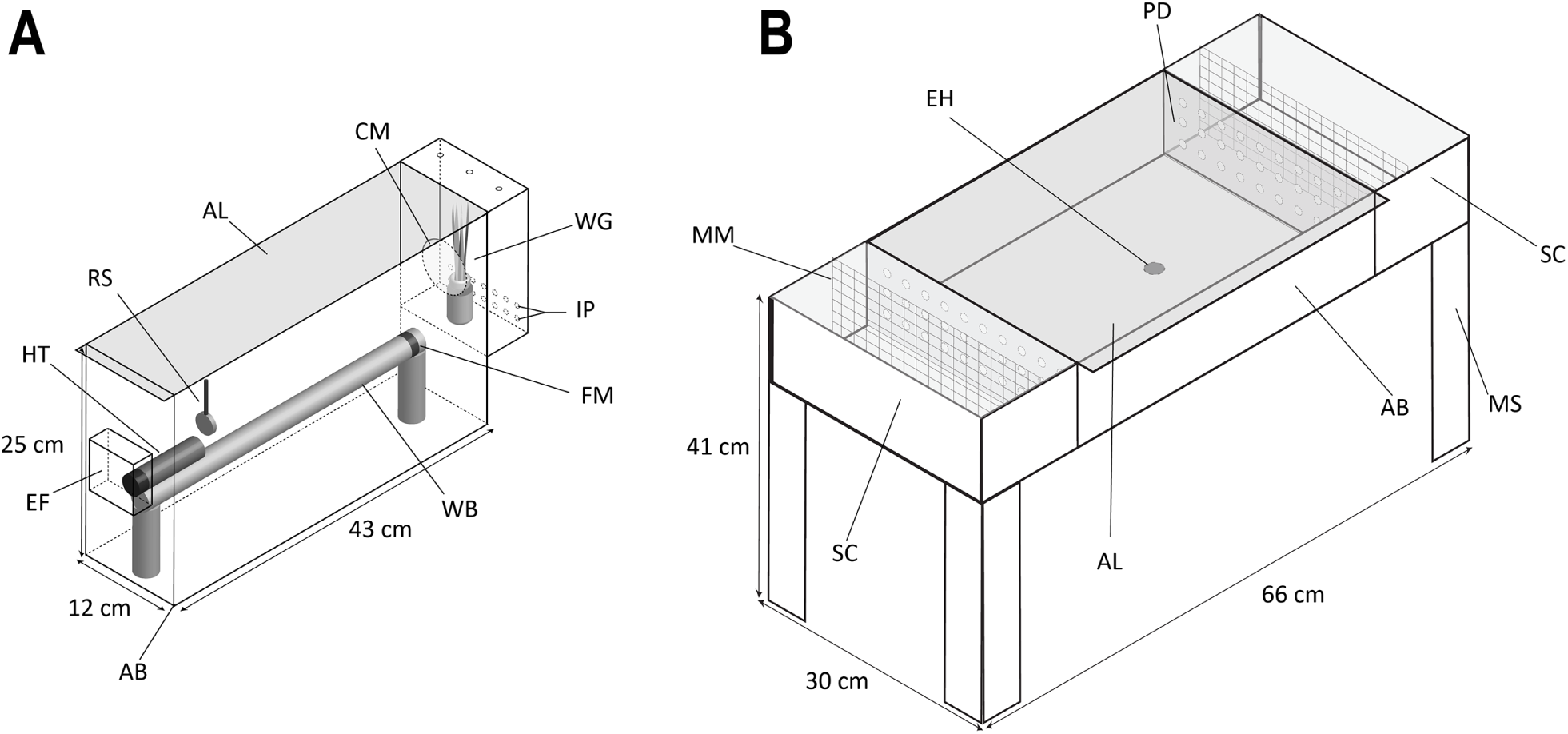
Diagrams of the two types of behavioural arena used. **A**: Diagram of beam-walk arena. AB = opaque acrylic box, AL = transparent acrylic lid, CM = fine cotton mesh, EF = air extract fan, FM = finish marker, HT = holding tube, IP = air intake perforations, RS = removable foam-board stopper, WB = wooden beam, WG = glass vial of wheat grass. **B**: Diagram of the Roessingh arena. AB = opaque acrylic box, AL = transparent acrylic lid, EH = entry hole, MM = metal mesh, MS = metal stand, PD = perforated transparent acrylic divider, SC = removable acrylic stimulus chambers.

#### Protocol

For testing, a locust was put head-first into an opaque holding tube fitted with a removable stopper at the head end. The rear end of the tube was plugged with a perforated polypropylene stopper. The holding tube was fastened with hook-and-loop fasteners to the top of the beam at the end nearest the fan, and the locust was given 10 min to acclimate. To start the trial, the stopper was removed, allowing the locust to emerge onto the beam and walk along it towards the source of the wheat odour. A 0.5 cm wide strip of black tape wrapped around the beam, 2 cm from the end nearest the wheat, marked the finish point. The trial ended after 10 min, irrespective of whether the locust had reached the finish point. The six arenas were recorded simultaneously at 30 frames per second and 640 × 480 pixels by a single Firewire video camera (Guppy F-033B, Allied Vision Technologies GmbH, Germany) mounted 105 cm above the arenas. This gave a spatial resolution of 9.56 pixels per cm.

#### Experimental Design

Beam-walk experiments used *N* = 59 solitarious and *N* = 50 gregarious Mauritanian strain locusts. Except during behavioural testing, locusts were kept at phase-appropriate densities throughout the experiment: individual rearing cages for solitarious locusts, and groups of 20–25 final instar nymphs in small cages (28 cm H × 10 cm W × 17 cm D) for gregarious locusts. On the day before the first trial, all food was removed from the cages. The next morning, at approximately 09:00, all locusts used in beam-walk assays were injected with 2.5 μl dimethyl sulfoxide (DMSO) per g body weight to also serve as vehicle controls for a separate pharmacological experiment not included here. DMSO is a widely used vehicle and is not expected to affect the two phases differently. One hour later, the locusts underwent their first 10 min arena trial, and were then returned to their holding cages for 3 h. The locusts underwent a second trial on the same day at approximately 14:00. After this, animals were once again returned to their holding cages and given food *ad libitum* for 2 h before being starved again overnight. The entire process was then repeated over the next 3 d, giving a total of *n* = 8 trials per animal (four in the morning and four in the afternoon). Using six arenas in parallel permitted testing of up to 30 animals over a 4 d period.

#### Behavioural Quantification

The recorded videos were used to measure the time between the opening of the tube and the moment when the whole of the animal’s head completely crossed the far edge of the finish marker. If the animal did not cross the mark within 10 min, the maximum distance it reached was recorded. Trials were discarded when the animal did not emerge from the tube, or left the beam before crossing it fully. For all valid trials, time taken to cross the beam or distance reached within 10 min were converted to a mean speed across the beam (*mean beam-walk speed*). Final sample sizes were 57 solitarious animals and 50 gregarious animals; two solitarious animals were removed as they only had one valid trial each. Power analyses are provided in the Supplementary Information.

### Roessingh arena behavioural assay

Experiments in the Roessingh arena used Leicester strain locusts. The Roessingh arena assay^[19,21]^ was originally developed for quantifying the behavioural phase state of individual locusts and has been used in many subsequent studies in *S. gregaria, Locusta migratoria* and *Chortoicetes terminifera* (See Cullen et al., 2017 for review ^[16]^).

The assay employs a rectangular central arena to contain the test locust, and removable stimulus chambers on each short end (Fig. 1B). Locusts were introduced into the arena through a 2 cm diameter entry hole in the centre of the floor. The side of each stimulus chamber facing the central arena was made of perforated transparent acrylic to allow both visual and olfactory stimuli to reach the test locust. One stimulus chamber held 30 gregarious final instar locusts as a crowd stimulus, whilst the other was empty. The positions of the stimulus and control sides were randomised for each session.

#### Protocol

To carry out the assay, a final instar test locust was placed into a 10 ml plastic syringe tube which was modified with a screw-lid attachment, screw-on cap, and covered with opaque tape. The test locust was left inside this tube to acclimate for 6 min. Following this, the tube was gently screwed onto the bottom of the entry hole, and the plunger gently raised to encourage the test locust upwards into the arena. Each test lasted for 7 min 45 s and were recorded by a FireWire video camera (Guppy F-033B, Allied Vision Technologies GmbH, Germany) mounted 70 cm above the arena. A metal mesh prevented stimulus animals from entering the camera’s field of view. The camera recorded at 30 frames per second and a resolution 640 × 480 pixels, giving a spatial resolution of 12.5 pixels per cm. The open-source software Swistrack (version 4.1^[22]^) was then used to track the x-y co-ordinates of the test locust in each video frame.

#### Experimental Design

We analysed 734 tracks from 324 solitarious individuals and 254 tracks from 81 gregarious individuals (Table 1) that were part of a large dataset collected over six years.

**Table 1:**
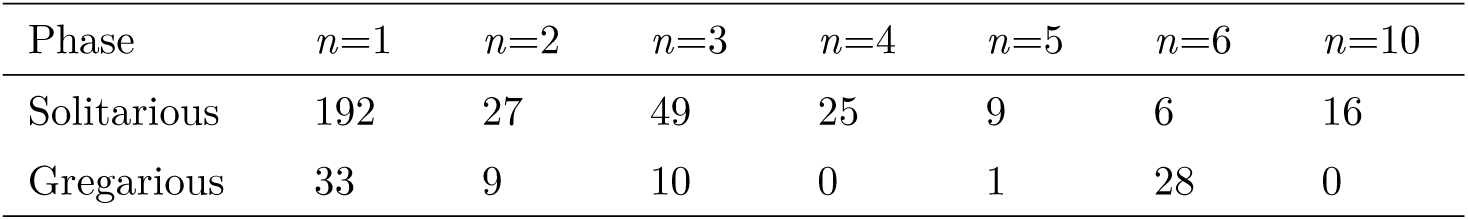
Number of tracks (n) per individual locust for each phase in the Roessingh arena dataset.

| Phase | $n=1$ | $n=2$ | $n=3$ | $n=4$ | $n=5$ | $n=6$ | $n=10$ |
| --- | --- | --- | --- | --- | --- | --- | --- |
| Solitary | 192 | 27 | 49 | 25 | 9 | 6 | 16 |
| Gregarious | 33 | 9 | 10 | 0 | 1 | 28 | 0 |

#### Behavioural Quantification: Mean Track-Speeds

The distance the animal had moved per frame was calculated from changes in x-y co-ordinates. These changes in position were then filtered using a rolling mean over a window of 11 frames to reduce apparent changes in position due to video pixel noise. Following this, mean speed was calculated by dividing total distance moved over the whole track by the track’s duration (*mean track-speed*). This measure corresponds to *mean beam-walk speed* in that it is based on the total assay duration including periods when the locust was stationary. Individuals with only one track were excluded from this analysis (Table 1). Note that *n*, the number of tracks (and hence *mean track-speed* observations) per individual was lower than the *n* = 8 in the beam-walk experiment, which reduces the accuracy of partitioning the total variance into between- and within-individual variance estimates. Power analyses are provided in the Supplementary Information.

#### Behavioural Quantification: Mean Bout-Speeds

Both the *mean beam-walk speed* and *mean track-speed* measurements include periods of hesitation in between movement bouts. To investigate variability over successive periods of continuous movement, we also analysed the mean speed of the individual walk-bouts (*mean bout-speed*). There were typically several walk-bouts per trial, defined as continuous periods during which the animal was consistently moving on the floor of the arena (i.e., not climbing) at a filtered speed greater than 1.35 cm/s and less than 12 cm/s for at least 18 frames out of every 20 (Fig. S1). Tracks from individuals for which only one track was available were included in this analysis (Table 1). This gave sample sizes of 3,653 bouts for solitarious individuals (median *n* = 6 per locust) and 5,034 bouts for gregarious individuals (median *n* = 58 per locust).

### Statistical Analyses

*Mean beam-walk speed*, Roessingh arena *mean track-speed* and Roessingh arena *mean walk-bout speed* were analysed in separate univariate log-normal mixed-effects models that are equivalent to Gaussian linear models with log-transformed movement speed as the response variable.

Distributions of speeds were greater than zero, positively skewed, and approximated well by a log-normal distribution (Fig. S2). If speeds *Y* are log-normally distributed, then *log Y* = *Y* * follows a normal distribution *N* with mean *μ* and variance *σ*^2^, which are also the parameters of the linked log-normal distribution, ℒ𝒩 thus

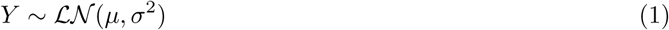

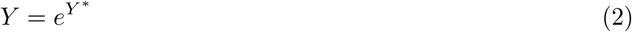

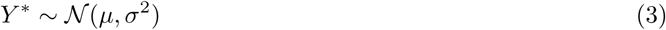

We use *μ* and *σ*^2^ when referring to means and variances on the log-scale specifically, to avoid confusion with means and variances on the data scale. As mean walk-bout speeds have a lower bound of 1.35 cm/s, this value was subtracted from mean walk-bout speeds to ensure the data match the support of the lognormal distribution (*Y* ∈ (0, + ∞)). This offset was added back onto *μ* after estimation, and does not affect *σ*^2^ (see Supplementary Methods for further information). The structure of all three models was

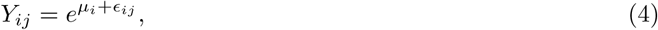

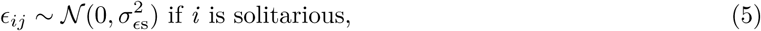

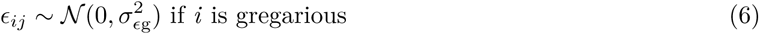

with *Y*_*ij*_ the speed of individual *i* in trial *j*; *μ*_*i*_ the mean log-speed of individual *i*; and _*ij*_ the deviation of individual *i*’s log-speed in trial *j* from its mean (*μ*_*i*_, residual error). These within-individual differences *ϵ*_*ij*_ were assumed to be normally distributed with mean 0 and variances 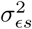 and 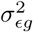 in solitarious and gregarious locusts respectively, thus allowing for phase-differences in within-individual variability. Each individual’s mean log-speed *μ*_*i*_ was in turn modelled as

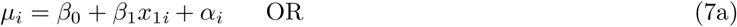

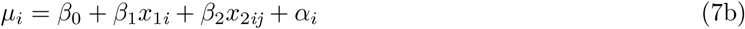

where Equation (7a) is the model fitted to the track-speed and bout-speed data, and Equation (7b) the model fitted to the beam-walk data. *β*_0_ is the group mean of the mean log-speeds of the solitarious individuals; *β*_1_ the difference between the solitarious and gregarious group means of the individual mean log-speeds; *x*_1*i*_ the phase of individual *i* (between-individual fixed effect; coded as *x*_1*i*_ = 0 for solitarious and *x*_1*i*_ = 1 for gregarious); and *β*_2_ the difference between the morning (a.m.) and afternoon (p.m.) group means of the individual mean log-speeds for both phases; *x*_2*ij*_ the time of day (a.m. or p.m.) for trial *j* of individual *i* (within-individual fixed effect, coded as *x*_2*ij*_ = 0 for a.m. trial and *x*_2*ij*_ = 1 for p.m. trial). Thus for Equation (7b) *β*_0_ and *β*_1_ refer specifically to the a.m. run.

For Equations (7a) and (7b), individual intercepts were modelled as

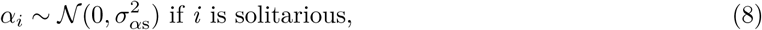

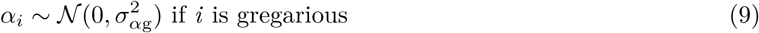

where *α*_*i*_ is the deviation of individual *i*’s mean log-speed from the group mean of means of the locusts of the same phase (random intercept). These between-individual differences *α*_*i*_ were assumed to be normally distributed with a mean of 0 and a variance that may differ between solitarious and gregarious locusts (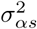 and 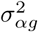, respectively), thus allowing for phase-differences in between-individual variability. Note that 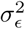 and 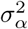 relate to the variances on the log-scale and not the observed variances on the data scale.

We use the geometric mean (GM) and geometric standard deviation (GSD) as measures of central tendency and dispersion of the log-normally distributed data. Motivations for this are discussed further in the Supplementary Methods. For a log-normal distribution,

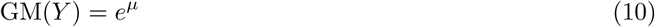

where *μ* is any group mean as estimated by the model. The geometric mean is also multiplicative such that, using the model in Equation 7a as an example, 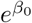 is the GM for the solitarious phase, 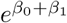 is the GM for the gregarious phase, and 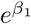 is the ratio of geometric means between the two phases. Therefore, we present differences between the two phases (and morning/afternoon runs for the beam-walk data) as the ratio of the difference in group means.

The GSD describes the interval containing 68.27% of the data as GM(*Y*) × GSD(*Y*)^±1^ and is calculated as *e*^*σ*^. In each locust phase *P*, we calculated separate estimates for within-individual GSD (GSD_*E*_) and between individual GSD (GSD_*α*_) as

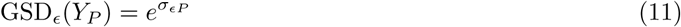

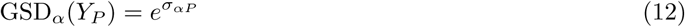

where *P* is either solitarious or gregarious.

All statistical analyses were carried out in the R statistical programming environment version 3.6.3^[23]^ within RStudio version 1.2.5003 ^[24]^. Models were fitted by Bayesian Markov Chain Monte Carlo (MCMC) estimation using the package brms ^[25,26]^. Priors used were only informative to the point that they restricted speed estimates to physically plausible values (see Supplementary Methods).

All terms estimated by the models are reported by giving the median of their posterior distribution along with the 95% credible interval (highest posterior density interval, HPDI; in the form of [lower, upper]). Distributions further from zero indicate increasing evidence for an effect. Calculations of GM and GSD were performed on posteriors of model estimates.

For all three models, chains and 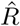^[27]^ indicated good convergence (Fig. S3) and all posteriors were normally distributed (Fig. S4). Our models assumed that residuals on the log scale (*ϵ*_*ij*_) were homoscedastic and normally distributed. Plots of the residuals verified that these assumptions were met for all three models (Fig. S5). Posterior predictive checks were used to verify that models fitted the observed data (Fig. S6).

## Results

We used two behavioural assays to quantify the locomotor behaviour of individual locusts. In the beam-walk assay, a food-deprived locust was presented with a source of food odour, which it could approach by walking upwind across a horizontal wooden beam. Since we standardised the satiation state of all locusts, the mean crossing speed served as a measure of a locust’s locomotor hesitation when approaching food. In the beam-walk arena, locusts were tested in the absence of conspecifics, regardless of their phase state. In contrast, the Roessingh arena assayed the locomotor behaviour of a single test locust in the presence of 30 conspecifics behind transparent perforated partitions. This combination of assays allowed us to separate reactive and intrinsic aspects of behavioural variability. We analysed (1) the mean speed of a locust’s track in the arena during the entire assay duration (including pauses in locomotion) as an overall measure of locomotor hesitation, and (2) the mean speed during each bout of continuous locomotion. Differences in mean crossing speed between the phases are presented as multiplicative differences with 95% HPD intervals (see Methods)

### Gregarious locusts show less locomotor hesitation

In both assays gregarious locusts moved less hesitantly than solitarious locusts. In line with the distributional assumptions of our log-normal models, the Bayesian estimates of geometric mean speeds closely matched the medians calculated from the raw data (Table 2). Further model validation is provided in the Supplementary Information.

**Table 2:**
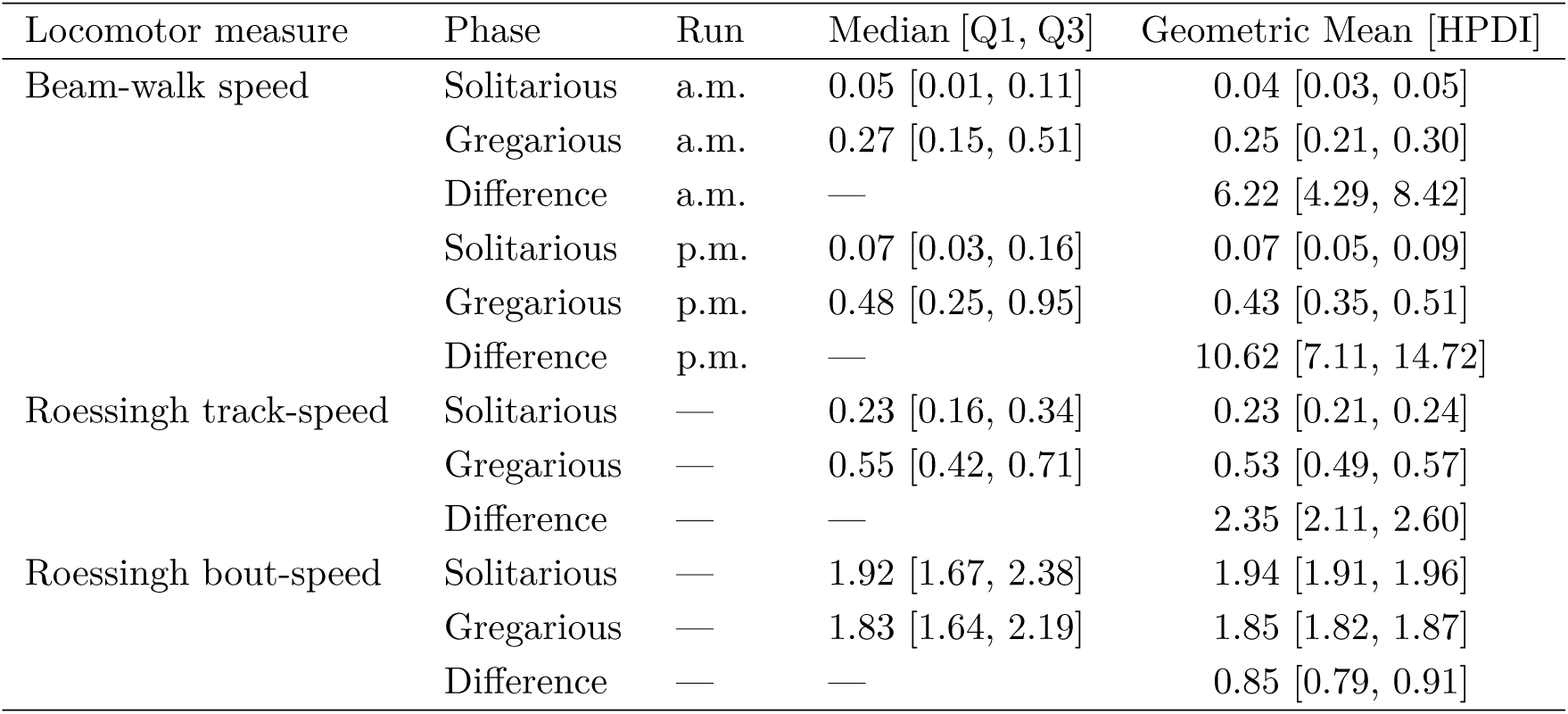
Comparison of raw group medians (Q1, Q3: first and third quartiles) and Bayesian mixed-effect model geometric mean estimates (HDPI: highest posterior density interval) for the three locomotor measures in solitarious and gregarious locusts. All speeds are in cm/s. Phase differences in geometric mean are presented as a fold difference.

In the beam-walk assay, gregarious animals crossed the beam 6.2 × [4.3, 8.4] faster than solitarious locusts in the morning (Fig. 2A), and 10.6 × [7.1, 14.7] faster in the afternoon (Fig. 2B). The track-speeds in the Roessingh arena were markedly higher than the beam-crossing-speeds, particularly for solitarious locusts (Table 2). Nevertheless, a clear phase difference in locomotor hesitation was apparent, as Roessingh arena track-speeds were 2.3× [2.1, 2.6] faster in gregarious locusts (Fig. 2C).

**Figure 2:**
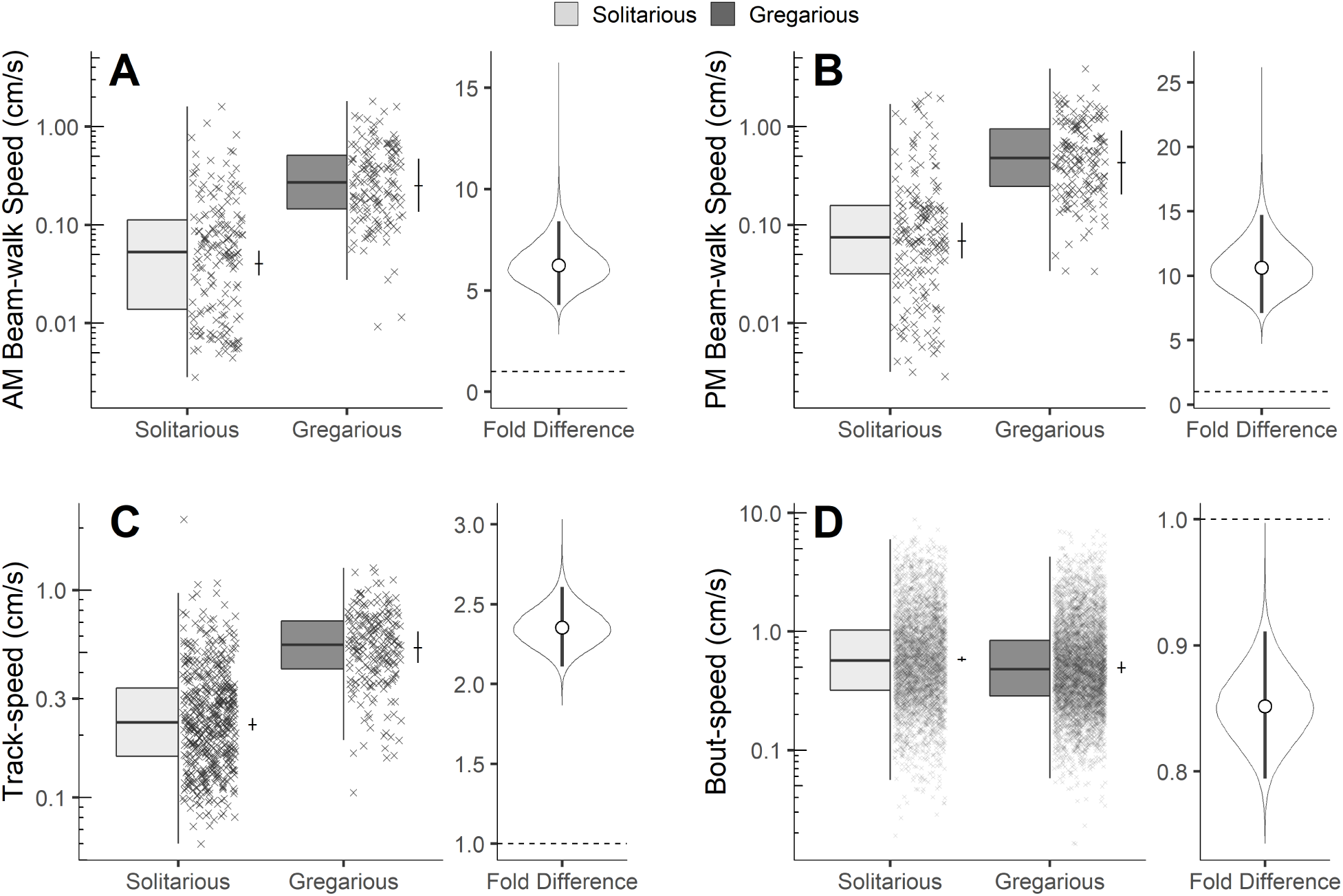
Gregarious locusts (dark grey) move less hesitantly than solitarious locusts (light grey) in both the beam-walk assay (higher beam-walk speeds; **A, B**) and the Roessingh arena assay (higher track-speeds, **C**), but their mean bout-speeds are almost identical (**D**). The box plots summarise the raw individual observations (×). The large crosses show Bayesian model estimates of geometric group mean (horizontal bars) and their 95% HPDI (vertical bars). Fold differences between the two phases are shown with separate axes in each panel. For these, violin shows the distribution of possible fold differences estimates from our Bayesian models, presented with median (white circle) and 95% HPDI (vertical bars). The dashed line indicates a fold difference of 1, i.e, no difference.

Roessingh arena bout-speeds were much higher than the track-speeds simply because they did not include locomotor pauses (Table 2). Remarkably, however, the mean speeds while walking for gregarious locusts were 0.85× [0.79, 0.91] those of solitarious locusts (Fig. 2D). The lower apparent locomotor speed of solitarious locusts therefore reflects a more intermittent mode of locomotion, rather than lower speed whilst moving.

### Gregarious locusts are less variable than expected

To compare the relative dispersion of the locomotor speeds between the two phases, we estimated their geometric standard deviations (GSD, Table 3), which are the multiplicative intervals around the geometric means that contain 66.7% of the observations (see Methods and Supplementary Information).

**Table 3:** Arithmetic standard deviations (SD) and Bayesian mixed-effect model estimates (HDPI: highest posterior density interval) of geometric standard deviations (GSD) for the three locomotor measures in solitarious and gregarious locusts. All speeds are in cm/s. Phase differences in GSD are presented as a fold difference.

| Locomotor measure | Phase | SD | Between-individual<br>GSD [HPDI] | Within-individual<br>GSD [HPDI] |
| --- | --- | --- | --- | --- |
| Beam-walk speed | Solitary | 0.29 | 2.67 [2.13, 3.34] | 2.78 [2.57, 3.02] |
|  | Gregarious | 0.47 | 1.73 [1.50, 2.02] | 2.07 [1.95, 2.21] |
|  | Difference | — | 0.65 [0.48, 0.83] | 0.75 [0.67, 0.82] |
| Roessingh track-speed | Solitary | 0.18 | 1.39 [1.30, 1.47] | 1.56 [1.51, 1.61] |
|  | Gregarious | 0.21 | 1.22 [1.13, 1.33] | 1.47 [1.41, 1.53] |
|  | Difference | — | 0.88 [0.80, 0.98] | 0.94 [0.90, 0.99] |
| Roessingh bout-speed | Solitary | 0.90 | 1.27 [1.20, 1.34] | 2.39 [2.34, 2.44] |
|  | Gregarious | 0.74 | 1.20 [1.15, 1.26] | 2.26 [2.23, 2.30] |
|  | Difference | — | 0.95 [0.88, 1.02] | 0.95 [0.92, 0.97] |

Compared with solitarious individuals, gregarious individuals were more similar to one another than expected from their means, in both measures of locomotor hesitation that we assessed. In the beam-walk paradigm, the between-individual GSD of the beam-walk speeds was 0.65 × [0.48, 0.84] lower in gregarious individuals (Fig. 3A). The difference was less pronounced, but in the same direction, for the Roessingh arena track-speeds, where relative between-individual GSD was 0.88 × [0.79, 0.98] lower in gregarious animals (Fig. 3C). Evidence for lower between-individual differences in the gregarious phase was weaker for the Roessingh arena bout-speeds, with an estimated 0.95× [0.88, 1.02] decrease in the GSD (Fig. 3E).

**Figure 3:**
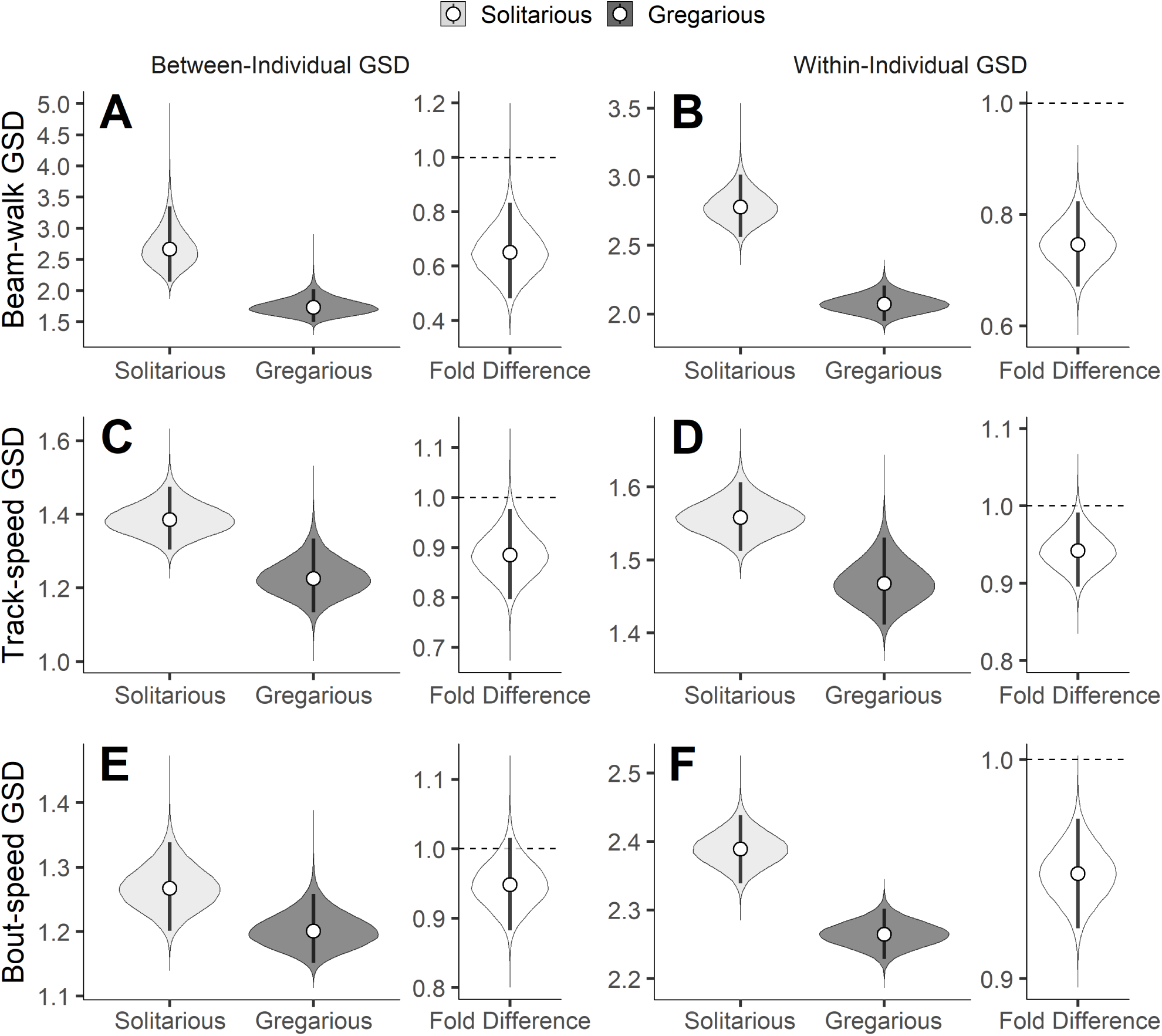
Gregarious locusts (dark grey) have locomotor speeds that are less variable relative to their means than solitarious locusts (light grey). Relative variability is measured by the geometric standard deviation (GSD), estimated separately between individuals (**A,C,E**) and within individuals (**B,D,F**). Distributions of posterior estimates are shown in density plots together with the medians (circles) and with the 95% HPDI (bars). **A,B**: GSD of beam-crossing speeds in the beam-walk arena. **C**–**F**: GSD of track speeds (**C,D**) and bout-speeds (**E,F**) in the Roessingh arena. Fold differences between the two phases are shown with separate axes in each panel. For these, the dashed line indicates a fold difference of 1, i.e, no difference.

Compared with solitarious individuals, the locomotor speeds of gregarious individuals were also more similar over repeated assays than expected given their means. Within-individual GSD was 0.75 × [0.67, 0.82] lower for the beam-walk speeds in gregarious individuals (Fig. 3B). Gregarious individuals were also more consistent in their Roessingh arena track-speeds and bout-speeds than solitarious individuals, but the differences were less pronounced: the within-individual GSDs of the track-speeds (Fig. 3D) and bout-speeds (Fig. 3F) were 0.94× [0.90, 0.99] and 0.95× [0.92, 0.97] lower, respectively, in gregarious individuals.

## Discussion

Polyphenisms evolve when contrasting selection pressures alternate too rapidly to permit fixation of one or the other phenotype ^[18]^. They therefore provide a window on evolutionary processes that can generate divergent stable phenotypes in environments with stable selection pressures ^[28]^. We have used the desert locust, a species that manifests two extreme phenotypes with contrasting predation ecologies, to examine the extent to which behavioural *variability* has evolved to be phenotypically plastic. Marked differences in *mean* behavioural phenotypes between isolated and crowded individuals are the defining feature of locusts, and a large body of work has documented that these differences outlast changes in the immediate social environment^[16]^. Our present study provides a step change from all previous work by asking whether density-dependent polyphenism in the desert locust extends to differences in the *variability* of behaviour. This has permitted us to test experimentally the important hypothesis ^[5]^ that different selection pressures on group-living and lone-living animals lead to adaptive differences in the variability of behaviour. No previous study has compared intrinsic variability of behaviour between two eco-phenotypes of the same species under tightly controlled and identical conditions.

For swarming animals, low behavioural variability should have a selective advantage ^[3]^, whereas for isolated animals, variable and unpredictable behaviour should be adaptive^[4,8]^. In line with these predictions, previous work in a variety of species has shown that individual animals adjust their behaviour to converge towards the group mean^[2]^. We have demonstrated that crowd-reared locusts are both more similar to one another in their behavioural means (lower relative between-individual variability) and more consistent over time (lower relative within-individual variability) than lone-reared locusts. We infer that contrasting selection pressures on a single genotype have led to the evolution of phenotypic plasticity in behavioural variability.

Reduced variability in large groups can be achieved by individuals following simple, instantaneously applied local rules, such as maintaining distance and alignment with neighbours, which lead to homogenous collective behaviour ^[7]^. In desert locusts, groups of gregarious individuals co-ordinate their collective behaviour by use of such local rules ^[29]^, but this cannot explain our results. Gregarious animals retained their lower relative variability even when tested in isolation in the beam-walk paradigm, and solitarious locusts retained their higher relative variability even in the presence of other locusts in the Roessingh arena. The different variabilities that we observed therefore do not result from instantaneous reactive responses to the presence or absence of conspecifics, but are intrinsic to the two phenotypes. The underlying regulatory mechanisms are therefore likely subject to selection, which, in more stable environments, would provide a substrate for the subsequent evolution of genetically determined differences in behavioural variability. Recent studies on yellow-bellied marmots and Trinidadian guppies provide evidence for genetic variance for behavioral predictability ^[15,30]^. As pointed out in these studies, demonstrating genetic variance for predictability in behaviours provides an important starting point for identifying both the environmental factors that genotypes respond to and the magnitude and direction of the evolutionary responses. We provide such empirical evidence that natural selection shapes individual predictability by demonstrating that locusts have evolved ‘variance polyphenism’. This polyphenism is likely adaptive as the direction of the phase difference aligns with predictions from ecological theory.

Solitarious locusts moved much more hesitantly in both locomotor assays. This superficially resembles previous reports that solitarious locusts walk more slowly than gregarious locusts in the Roessingh arena^[16,31,32]^, but we find that the speed when walking (‘walk bout speed’) is very similar in the two phases — and in fact is slightly lower in gregarious locusts.The discrepancy arises because previous studies scored ‘walk speed’ as zero for locusts that did not walk^[31]^. This underestimates the walk speed of solitarious locusts because these animals account for the majority of assays where walking did not occur. The more hesitant locomotion of solitarious locusts arises not from slower but from more intermittent walking. Such intermittent locomotion is associated with predator avoidance through behavioural crypsis in other species ^[33]^ and is likely to do so in solitarious locusts. Gregarious locusts, however, have shorter^[34]^ and less variable^[35]^ pauses when moving in a crowd, which facilitates their collective motion important in swarm coherence^[29]^. The lower intrinsic variability of behaviour we identified in gregarious locusts will work in tandem with instantaneous local interactions to create homogeneous collective behaviour.

Our finding of reduced between-individual differences in gregarious locusts is in direct contrast with the ‘social niche specialisation’ hypothesis, which posits that social interactions amplify between-individual differences^[36]^. Evidence to support this hypothesis has been found in some social mammals^[37]^ and invertebrates such as crickets^[38]^ and spiders^[39]^, yet in other social animals, group living is associated with social conformity ^[40]^. Neither hypothesis, however, provides a useful framework for explaining behavioural differences among individuals in animals such as the desert locust, which are not social beyond simple aggregation. Thus, the contrast between our findings in locusts and those in other non-social insects emphasises that the effect of group-living on between-individual variability reflects the ecology of the species in question.

Prior empirical work on protean anti-predator behaviour is both limited and conflicted. Hermit crabs increase within-individual variability in the presence of a predator ^[8]^, and variability of prey behaviour influences both hunting success in jumping spiders^[41]^ and the success with which humans capture simulated prey ^[42,43]^. In contrast, however, tadpoles that experienced predator cues during ontogeny showed decreased within-individual variability in the presence of predators ^[12]^. Our results are not able to directly resolve this conflict, as we did not test the effect of predator cues on behavioural variability. Although phase change leads to protean behaviour in solitarious locusts, the increased variability is driven proximately by a history of isolation from conspecifics, rather than by cues from predators.

In contrast to most previous related work (e.g. Prentice et al. 2020 ^[15]^; Briffa, 2013 ^[8]^) our models measured individual variances *relative to their corresponding mean*. This explicitly acknowledges the distributional properties of the observed data, which are well approximated by a log-normal distribution, in that they have a lower bound of 0 and, importantly, variances that scale with the mean. This reflects the trivial fact that a low mean necessarily implies a lower variance if the distribution has a lower bound of zero; variance can only increase through larger data points, which result in a higher mean. Consequently, gregarious locusts - which are more active - are also more variable in absolute terms than solitarious locusts. Our data share this with many biological variables^[44]^. For example, higher temperatures cause both higher means and more variability in exploratory behaviour in crickets. In this case temperatures also change the coefficients of variation (CV) as estimated from a Gaussian model ^[13]^. In contrast, log-normal models directly account for the means-variance relationship and, if appropriate, permit explicit analysis of that relationship, as quantified by the geometric standard deviation (GSD). In this light, any difference in variance that is not reflected in a difference in GSD indicates uninteresting distributional constraints rather than important biological ones (e.g., genetic constraints ^[45]^).

To conclude, we present clear evidence for differences in behavioural variability that outlast the social conditions which trigger them. This polyphenism in behavioural variability implies a mechanism regulating behavioural variability that evolved in response to conflicting selection pressures. Our findings provide empirical evidence linking differences in group living to the evolution of differences in behavioural variability.

## Acknowledgements

We thank Ben Warren and Brendan O’Connor for comments on the manuscript. We also thank Chanida Fung, Anthony Vencatasamy, Jake Cranston and Rubèn Gonzalez-Colom for assistance with locust husbandry. This work was funded by BBSRC Research Grant BB/L02389X/1 awarded to SRO and TM, a BBSRC MIBTP studentship awarded to JMS and a studentship from the College of Life Sciences (University of Leicester) to BC.

## Author contributions

BC, TM and SRO conceived the study. BC collected beam-walk data. JMS collected Roessingh arena data. BC and SRO analysed data. BC, TM and SRO wrote the manuscript.

## Supplementary Information

### Locust Rearing

Gregarious locust cages were kept under a 12 h:12 h light-dark cycle, with temperatures of 35–38°C during the day and 25°C during the night. In order to generate solitarious locusts, gregarious animals were isolated as hatchlings and reared to adulthood in isolation in a separate facility to serve as breeders. The offspring of these breeders were then used as experimental animals.

Animals were initially isolated in small transparent plastic pots (8 cm H × 5 cm diameter). Each pot was provided with an independent clean air supply drawn from outside the building and heated to 36°C (day) or 25°C (night) and 20–25% relative humidity. The solitarious facility followed the same day–night cycle as the gregarious facility. Pots contained a bundle of fresh wheat seedling leaves, and an absorbent string wick. These dipped into a tub of water (7 cm H × 5 cm diameter) which sat below the pot to keep the wheat fresh, and allow the hatchling to drink via the wick. Wheat bran flakes were sprinkled over the floor of the upper pot.

Around the 3^*rd*^ or 4^*th*^ instar, animals were transferred to larger metal cages (10 cm L × 10 cm W × 20 cm H). These cages contained a vial of wheat grass seedling leaves and a dish of wheat bran which were replaced twice a week. Heated outside air was pumped into the cages with 45 air changes per hour. Solitarious animals could thus be reared in large numbers but in complete isolation from the sight, smell and touch of conspecifics.

### Beam-walk Assay

#### Arena Design

A rectangular enclosure (43 cm L × 12 cm W × 24 cm H) was constructed from matte white acrylic walls and floor with a removable transparent acrylic lid. Inside, a 38 cm long wooden dowel rod of 2.5 cm diameter was raised on dowel stilts 11 cm above the centre of the arena floor. A circular opening (6 cm diameter; centre point 14 cm above floor level) was cut into each of the narrow end walls. A perforated box containing a vial of fresh wheat grass seedling leaves was fitted behind one opening, which was screened with cotton fabric. The opposite opening was fitted with a small extraction fan (12 V, 0.14 A; model AGE06025B12M, Crown, Taipei, Taiwan) to continuously draw room air with wheat grass odour through the arena. The bank of six arenas was lit from above with three equally spaced desk lamps, each centred on two arenas, resulting in an illumination of 1,500–2,000 Lux.

Holding tubes for loading animals into the arena were 2.2 cm inner diameter, 7.7 cm long; cut from 35 ml tubes, (Sarstedt AG & Co. KG, Germany, cat. no. 58.537). Polypropelyne stopper was 1.5 cm long 2.2 cm diameter (Sarstedt, cat. no. 65.790) and was drilled to allow air flow through the tube.

#### Video Capture

The six arenas were recorded simultaneously at 30 frames per second and 640 × 480 pixel resolution using a single Firewire video camera (Guppy F-033B, Allied Vision Technologies GmbH, Germany) fitted with a 1/1.8” 4.4–11 mm high resolution varifocal lens (Edmund Optics, Ltd, UK; stock #68-665). The camera was mounted 105 cm above the beam and connected to an Intel Core 2 Duo E8400 PC (2 × 3.0 GHz) running SUSE Linux Enterprise Desktop 12 SP2. Coriander video capture software (version 2.0.2; https://sourceforge.net/projects/coriander/) was used to control the camera. Videos were initially saved as uncompressed raw files and later MPEG-4 compressed using FFmpeg software (version 3.4.6; https://ffmpeg.org).

### Roessingh Arena

#### Arena Design

The apparatus consisted of a central white acrylic arena with a clear acrylic lid (40.5 cm L × 28.7 cm W × 10.0 cm H) to contain the test locust and removable stimulus chambers on each end (28.3 cm L × 13.0 cm W × 10.0 cm H). The central chamber contained a 2 cm diameter entry hole for the test locust in the middle of the arena floor. To reduce the frequency of locusts climbing on the lid, 3 mm thick white Teflon sheets were glued to the long inner walls. The side of each stimulus chamber facing the central chamber was made of perforated transparent plastic to allow both visual and olfactory stimuli to reach the test locust. One stimulus chamber held 30 gregarious 5^*th*^ instar locusts as a crowd stimulus, whilst the other was kept empty. The positions of the stimulus and control sides were randomised for each session. To avoid detection of the stimulus group by the tracking software, a metal mesh was mounted vertically within each stimulus chamber, leaving a space of 4.5 cm between the stimulus group and the transparent perforated acrylic front. The arena floor was covered with an A3 sheet of white paper that was changed at the beginning of each experimental day. The arena was mounted in a metal frame that was covered with white cotton sheets and illuminated symmetrically by placing a flicker-free LED strip light (DFx Planetsaver LED Ultra Low Profile, 12V dc; 5000K Pure White, 16 × 16 × 320 mm) on top of each stimulus chamber. The syringe tube used to insert animals into the arena was 2 cm diameter and 8 cm length

#### Video Capture

A FireWire video camera (Guppy F-033B, Allied Vision Technologies GmbH, Germany) fitted with a 1/3” 3.4–10.5 mm high resolution varifocal lens (Computar LLC, United States, model T3Z3510CS) was mounted 70 cm above the arena. Videos were captured and converted to MP4 files as described in the ‘Beam-walk Assay’ section.

**Figure S1:**
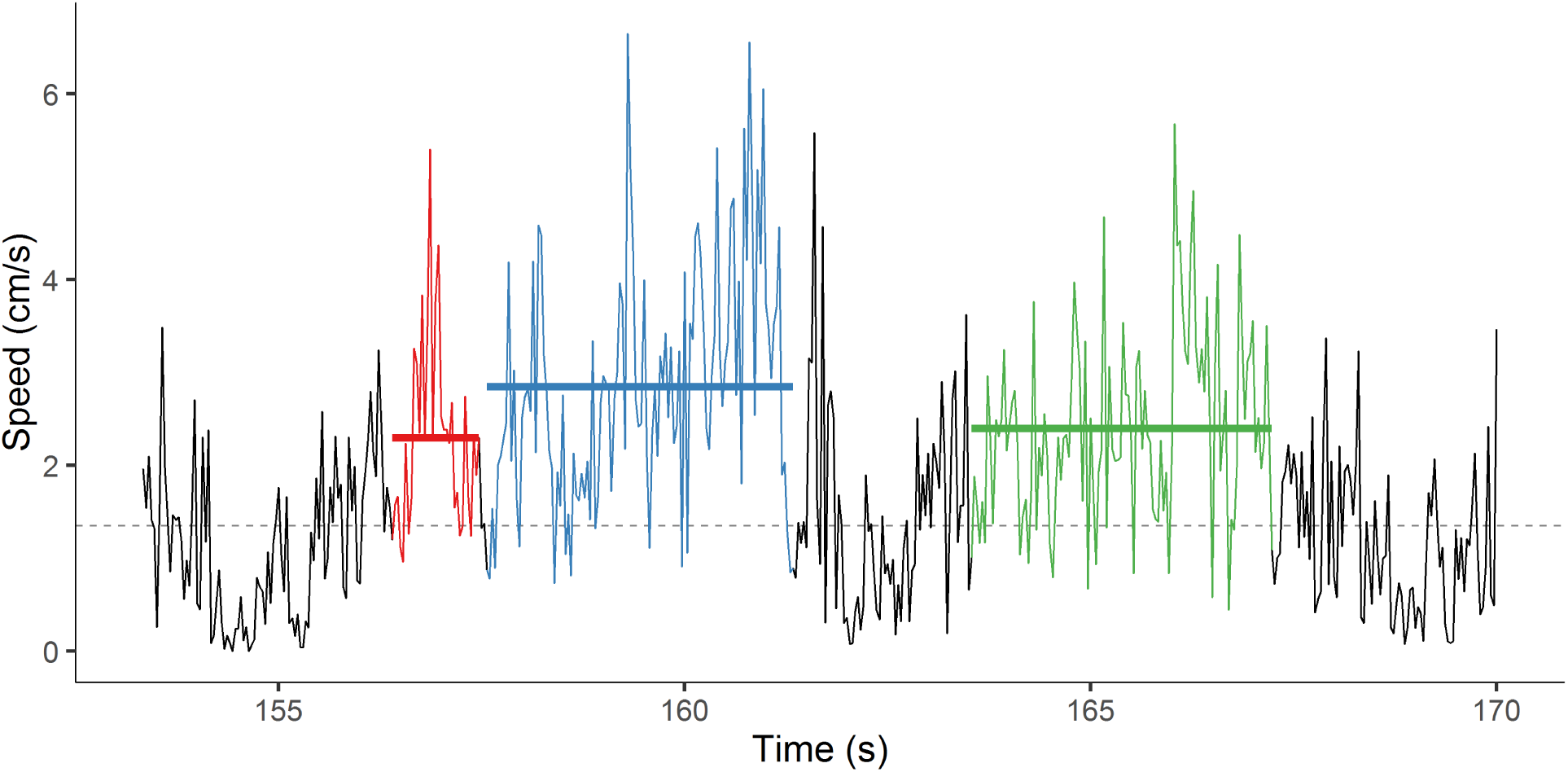
Example of how mean walking bout speeds were derived from a locust’s instantaneous speed as calculated from the frame-to-frame displacement in the movement trajectory. The dashed horizontal line represents the threshold above which the animal was considered as walking. The three coloured sections are time regions during which the instantaneous speed was above the threshold for at least 18 out of each 20 consecutive frames: these are defined as ‘walking bouts.’ The solid horizontal lines show the mean movement speed (y-axis) for each bout.

### Statistical Analysis

#### Geometric means

In our log-normal models, the *β*-coefficients refer to differences in the group means *μ*_*G*_ of the log-transformed speeds *Y*. For each group *G* (solitarious, gregarious; separately for a.m. and p.m. trials in the beam-walk assay), we calculated an estimate for *μ*_*G*_ from the model estimates of the *β*-coefficients. Estimates of the group *geometric means* GM(*Y*_*G*_) of the speeds were obtained by exponentiation of the estimates for *μ*_*G*_, since GM(*Y*) = *e*^*μ*^ for the log-normal distribution. Unlike the arithmetic mean, the geometric mean is independent of *σ*^2^ and coincides with 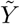, the median of *Y*. The geometric mean was therefore our preferred measure of central tendency.

#### Geometric Standard Deviations

Unlike the group means and group variances, the within- and between-individual variance components on the data scale are not meaningfully defined for a log-normal mixed-effects model, because for a given value of *σ*^2^, the variance of the log-normal distribution depends on *μ*. Although all locusts of one phase have the same within-individual *σ*^2^ (as per model assumption), individual locusts of one phase have different individual mean log-speeds *μ*_*i*_, and thus different within-individual variances. Similarly, if the two phases differ in the group means of their log-speeds, a given value of 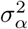 would manifest in different between-individual variances.

An alternative measure of dispersion on the data scale that is independent of *μ* is provided by the (multiplicative) geometric standard deviation, GSD(*Y*) = *e*^*σ*^. The GSD is a unit-less factor that describes the dispersion around the geometric data mean: the interval GM(*Y*) × GSD(*Y*)^±1^ covers 68.27% of the data. In each locust phase *P*, we calculated separate estimates for the within-individual 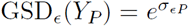 and for the between-individual 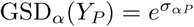.

**Figure S2:**
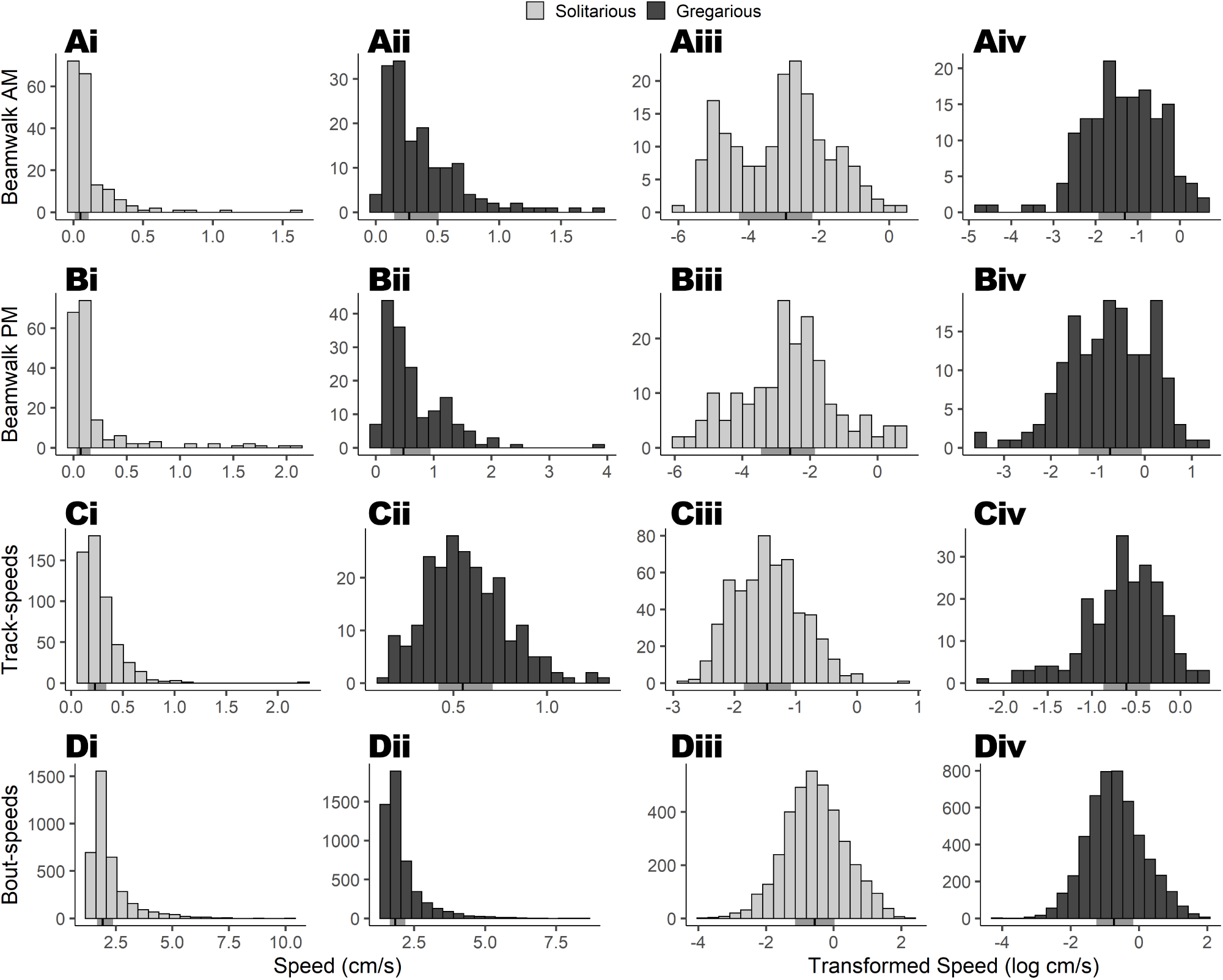
Histograms showing distributions of raw (i, ii) and log-transformed (iii, iv) speeds from the beam-walk in the morning (A) and afternoon (B), the Roessingh arena track-speeds (C) and bout-speeds (D). Solitarious speeds (light gray) are shown in i and iii, and gregarious speeds (dark gray) are shown in ii and iv. The vertical black line below each histogram is the median, and the shaded interval around it is the interquartile range.

**Table S1:** Prior distributions of fixed effects in all three models. *β*_0_ is the intercept, and represents the mean speed of solitarious animals (in the morning for the beam-walk model). *β*_1_ is the difference in means between a solitarious and gregarious animal (in the morning for the beam-walk model). *β*_2_ only appears in the beam walk-model and is the difference between the morning and afternoon mean speeds in both phases.

| Model parameter | Mean ($\mu$) | Standard deviation ($\sigma$) |
| --- | --- | --- |
| $\beta_0$ | 0 | 2 |
| $\beta_1$ | 2 | 6 |
| $\beta_2$ | 1 | 2 |

#### Three-parameter log-normal distribution

Since mean bout-speeds had a lower bound of 1.35 cm/s due to their generation process, they did not match the support of the log-normal distribution, *Y* ∈ (0, + ∞). Thus, we modelled mean bout-speeds using a three-parameter log-normal distribution. This shares the parameters of the mean *σ* and standard deviation *σ* with a standard log-normal distribution, but also features the threshold parameter *λ*. Therefore, for a log-normal distribution

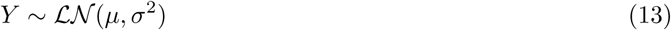

the corresponding three-parameter version is

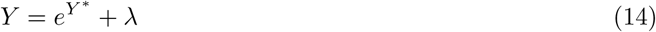

For our bout-speed model, this simply meant that 1.35 was subtracted from speeds before log-normal modelling occurred (to ensure that support was met), then 1.35 was added on again when model estimates of GM were calculated on the data scale, that is

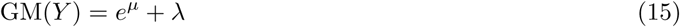

As the GSD is a measure of dispersion around the GM, independent of the value of the GM, it is likewise unaffected by the presence or value of the threshold parameter *λ*.

#### MCMC Model Fitting

Each model was fitted by running eight parallel MCMC chains for 125,000 iterations following a warmup of 10,000 iterations. Only every 10^*th*^ sample was retained, giving 100,000 total iterations across all 8 chains. Convergence was assessed visually and using the Gelman-Rubin convergence diagnostic 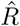^[27]^. This is a ratio of the variance within chains to the variance between chains, and is ideally 1 for a converged model.

Priors for fixed effects were chosen to restrict model estimates to physically plausible values. These priors were normal distributions with means and standard deviations shown in Table S1. Random effects all used a weakly informative half-t prior (*v* = 3, *μ* = 0, *σ* = 10).

**Figure S3:**
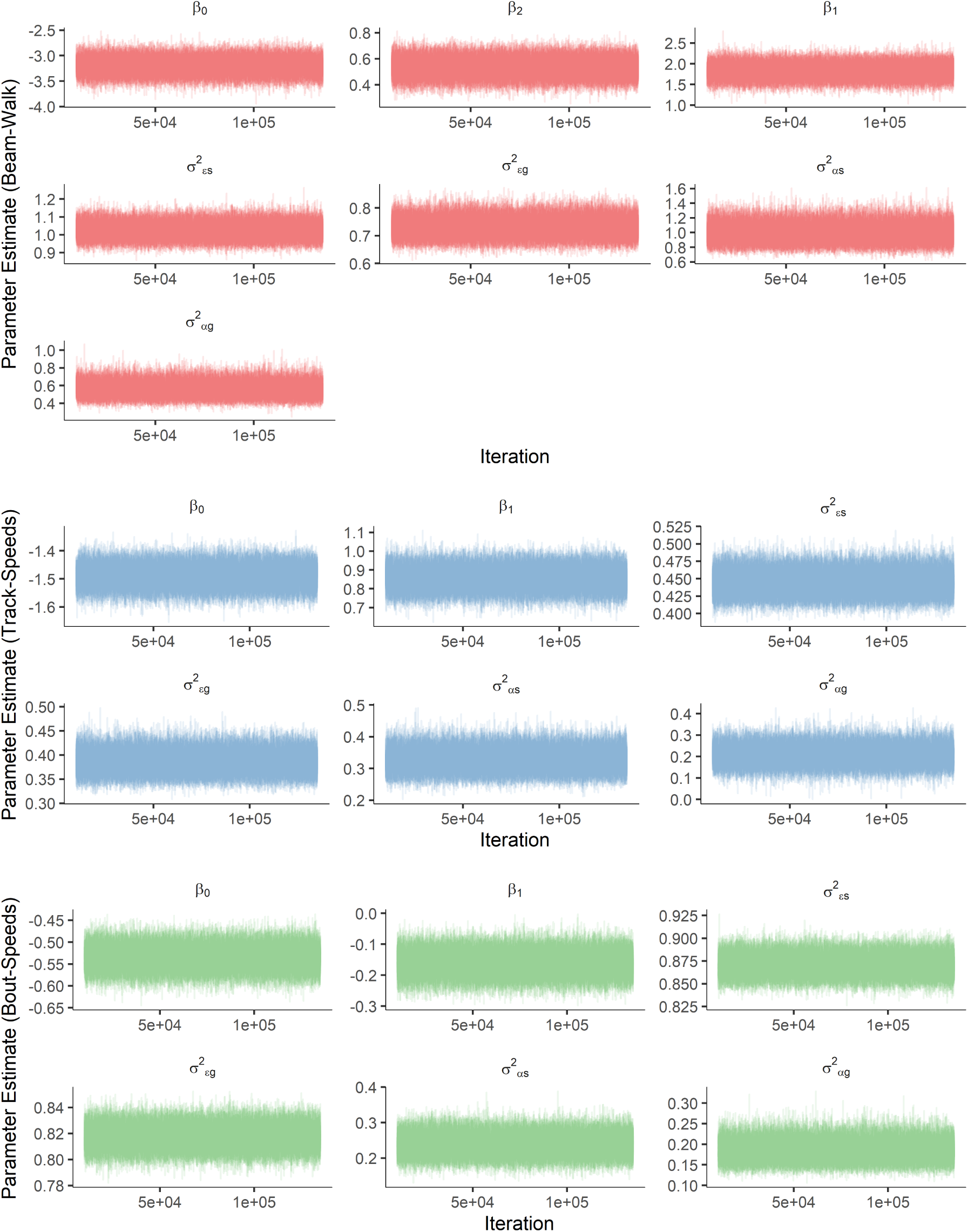
Sampling behaviour of the chains for each model. Beam-walk model parameters are in red, track-speeds in blue and bout-speeds in green. The absence of any discernible patterns between chains or over iterations indicates good convergence.

**Figure S4:**
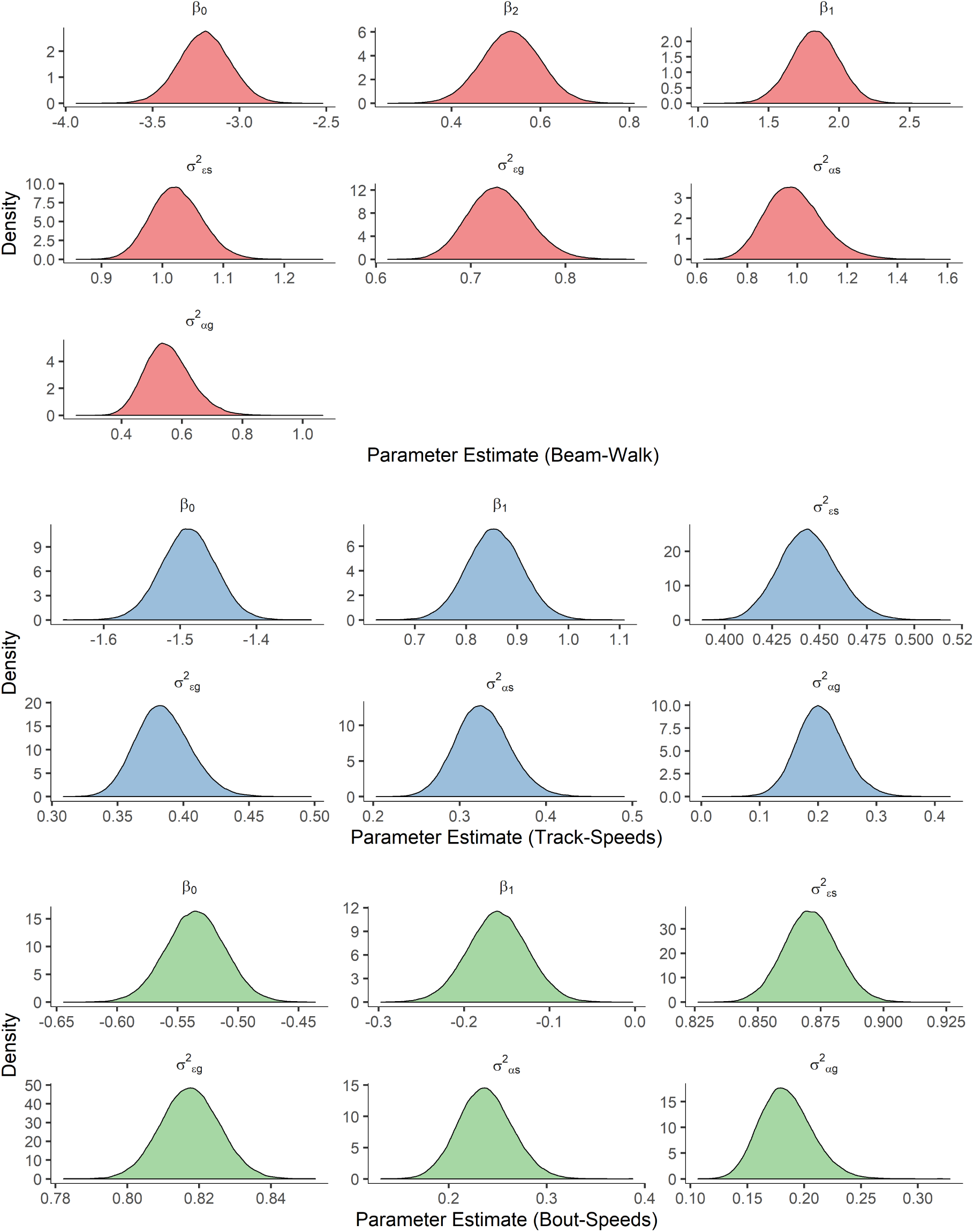
Posteriors for each parameter estimated by the models. Beam-walk model parameters are in red, track-speeds in blue and bout-speeds in green. Posterior distributions here are very close to normal, indicating good convergence.

**Figure S5:**
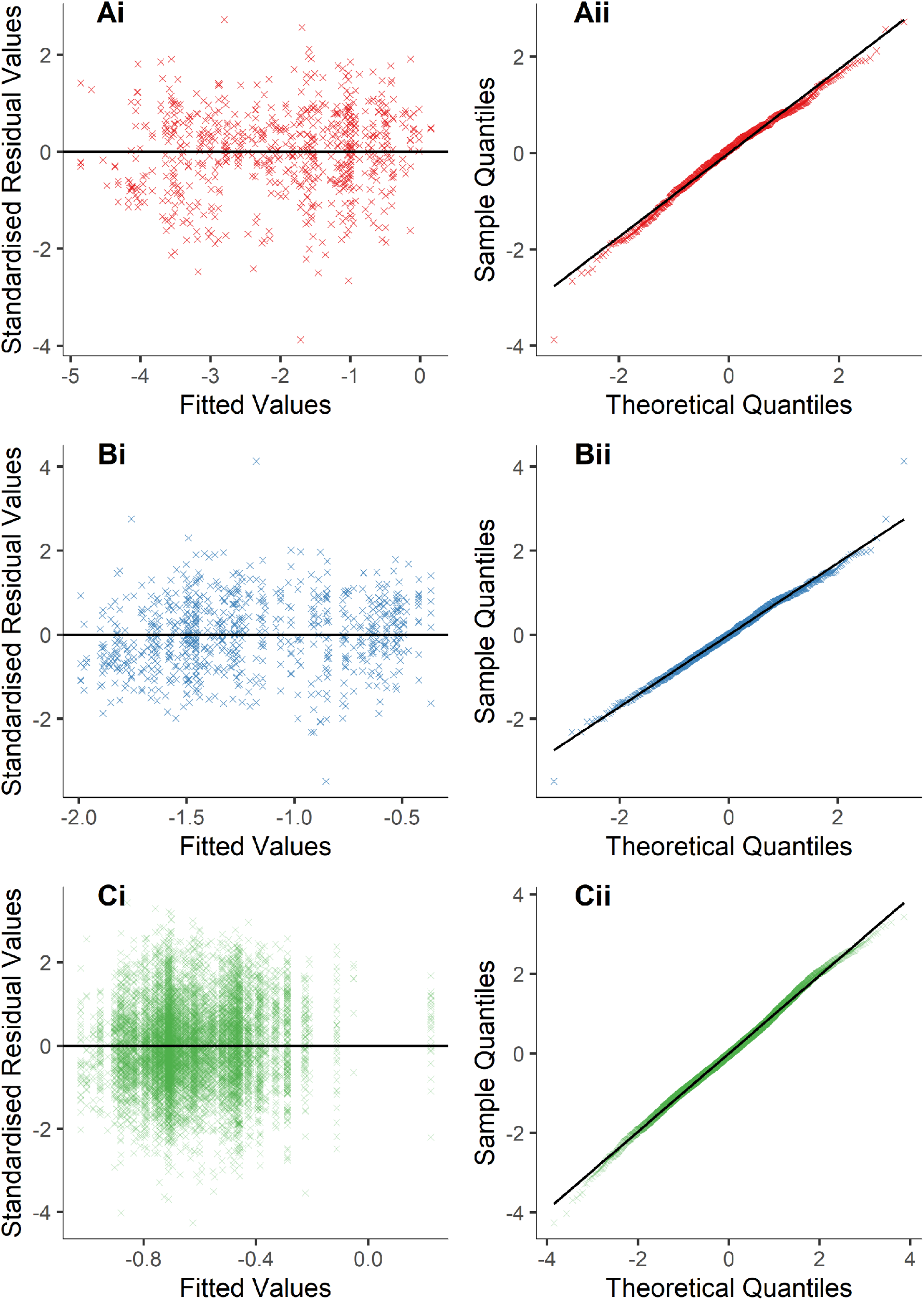
Verification of model assumptions. A) Assumptions for beamwalk model. B) Assumptions for track-speeds model. C) Assumptions for bout-speeds model. Ai, Bi and Ci show residuals vs fitted values, which should show a similar normal distribution around zero along the range of x axis values if assumptions are met. Aii, Bii and Cii show Quantile-Quantile plots, where all residuals should fall on or close to the line if assumptions are met.

**Figure S6:**
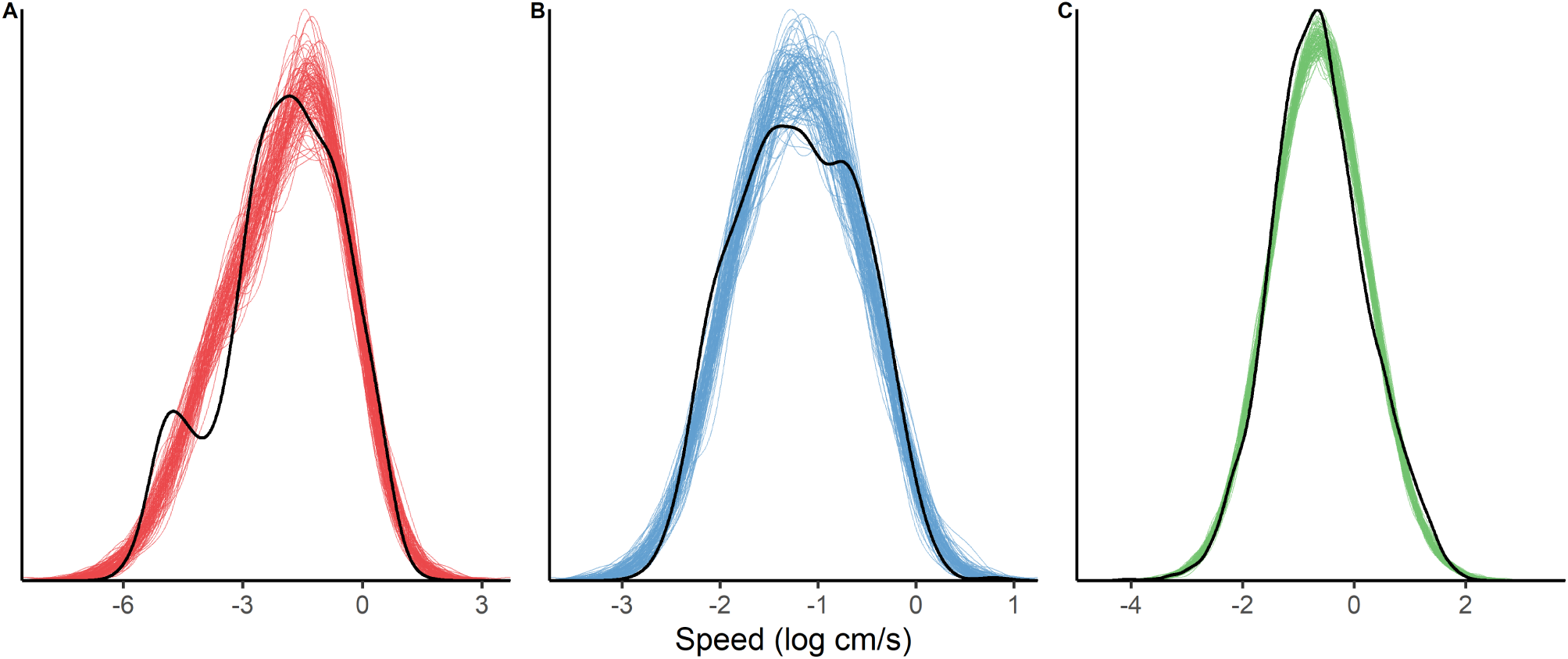
Posterior predictive checks for the beam-walk model (A), the track-speed model (B) and the bout-speeds model (C). For each model, the black line represents the distribution of the log-transformed speeds in the raw data. An estimate of each parameter was then taken from the model posteriors and used to simulate speed values. This process was repeated 100 times, and the distributions of each resulting simulation are shown in the overlaid coloured lines. The black line follows these distributions well, which suggests that the model is a good representation of the process that generated the observed data. Slight discontinuities reflect where data slightly deviate from an ideal normal distribution.

### Power Analysis

A power analysis was carried out to determine how robustly our statistical approach could detect differences in geometric standard deviation (GSD) at the sample sizes we used (number of individuals and number of measurements per individual). Power calculations for these types of models are non-trivial: We therefore simulated a large dataset with a known difference in GSD, drew repeated subsets from that larger dataset, and determined the proportion of subsets in which an effect was detectable. This process was then iterated over a range of effect sizes.

#### Effect Size Specification

As discussed in the main paper, our mixed-effects models split variability into two components: between-individual (*α*) and within-individual (*ϵ*). We first generated vectors of all the *α* and values that we wished to profile. These were expressed as fold increases in the ‘solitarious’ GSD based on a constant ‘gregarious’ GSD (which we took from our own data for each assay).

Since it is likely that the ability to detect a difference in one of these components depends upon the magnitude of both components (i.e, total variance), we tested all combinations of the specified fold differences in *α* and. Any one combination of *α* and was therefore an effect size (*E*) and was iterated on as the outer loop of our simulation.

#### Dataset Simulation

Estimates of fixed effects (phase, and run for the beam-walk data) were taken from the models used in the main paper. GSD values were created by using the GSD_*α*_ and GSD_*ϵ*_ estimates for the gregarious phase from the final models, then multiplying each by the fold differences specified as the effect size for *α* and *ϵ*.

We then used these parameters to simulate a dataset. For each individual in our dataset, we first drew an individual intercept which represented between-individual variance (*α*). This was drawn from a normal distribution with a mean of 0 and variance of log(GSD_*α*_) based on the individual’s phase.

Next, we drew the correct number of ‘trials’ for the individual from a log-normal distribution with a phase-specific mean plus the individual intercept, and a phase-specific variance of log(GSD_*ϵ*_). For the beam-walk simulation, the ‘run’ fixed effect was also added onto the mean for half of the trials. This process was then repeated for each individual until a full dataset is created.

#### Modelling

A model was then run with the same structure, priors and parameters discussed in the main paper. From this model, we stored the median ± HDI for each parameter, as well as the median ± HDI for the difference in posteriors of GSD_*α*_ and GSD _*ϵ*_ for each phase. A dataset was recorded as ‘detecting a difference’ if the 95% HDI of this posterior did not overlap with 0.

A new dataset was then simulated, and this process was repeated for a total of 100 datasets per combination of effect sizes. Table S2 shows the combinations of effect sizes profiled for each dataset, and Figure S7 summarises the simulation process as a flowchart.

## Results

Figure S8 shows the power curves for different effect sizes for each parameter. Generally speaking, a larger GSD_*ϵ*_ increases power requirements to detect the same difference in GSD_*α*_. As such, for each of GSD_*ϵ*_ and GSD_*α*_ we present the minimum detectable effect sizes (at 0.8 power) here as a mean ± sd calculated across all profiled values of the other. For ease of comparison with fold differences shown in the main paper, we present these differences as a fold decrease in the ‘gregarious’ group, relative to the ‘solitarious’ group. Lower values therefore indicate a bigger difference, so a minimum detectable difference closer to 1 indicates higher power.

For the beam-walk dataset, our minimum detectable effect sizes were a difference in GSD_*ϵ*_ of 0.905 ± 0.004 and a difference in GSD_*α*_ of 0.744 ± 0.015. For the track-speed dataset, our minimum detectable effect sizes were a difference in GSD_*ϵ*_ of 0.933 ± 0.002 and a difference in GSD_*α*_ of 0.879 ± 0.009. For the bout-speed dataset, our minimum detectable effect sizes were a difference in GSD_*ϵ*_ of 0.969 ± 0.002 and a difference in GSD_*α*_ of 0.922 ± 0.003.

**Figure S7:**
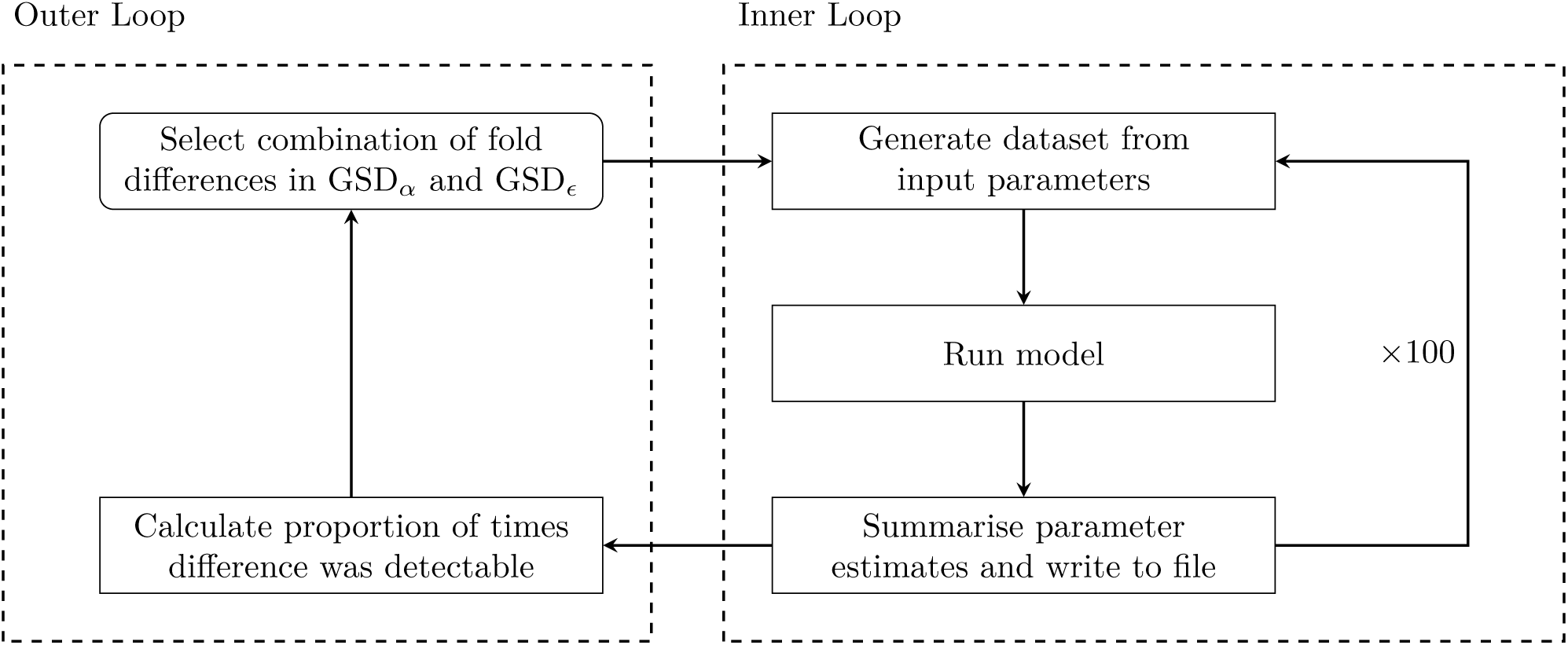
Flowchart summarising our power analysis approach. For each dataset, the outer loop was performed for all combinations of GSD_*ϵ*_ and GSD_*α*_ found in Table S2, and was executed sequentially. The inner loop was executed in parallel, with 20 iterations being performed simultaneously by an Intel Xeon Gold 24-core processor.

**Table S2:** Percentage increases in between-individual (GSD_*α*_) and within-individual (GSD_*ϵ*_) dispersion profiled by our power analysis. For each model, every possible combination of GSD_*α*_ and GSD_*ϵ*_ was profiled.

| Model | $\text{GSD}_\alpha$ values | $\text{GSD}_\epsilon$ values |
| --- | --- | --- |
| Beam-walk | 0, 5, 25, 35, 50, 100 | 0, 5, 10, 15, 25, 50, 75, 100 |
| Track-speeds | 0, 7.5, 10, 12.5, 15, 20, 25 | 0, 2.5, 5, 6, 7.5, 12.5, 15, 25 |
| Bout-speeds | 0, 2.5, 5, 7.5, 10, 15 | 0, 1, 2, 3, 4, 5 |

**Figure S8:**
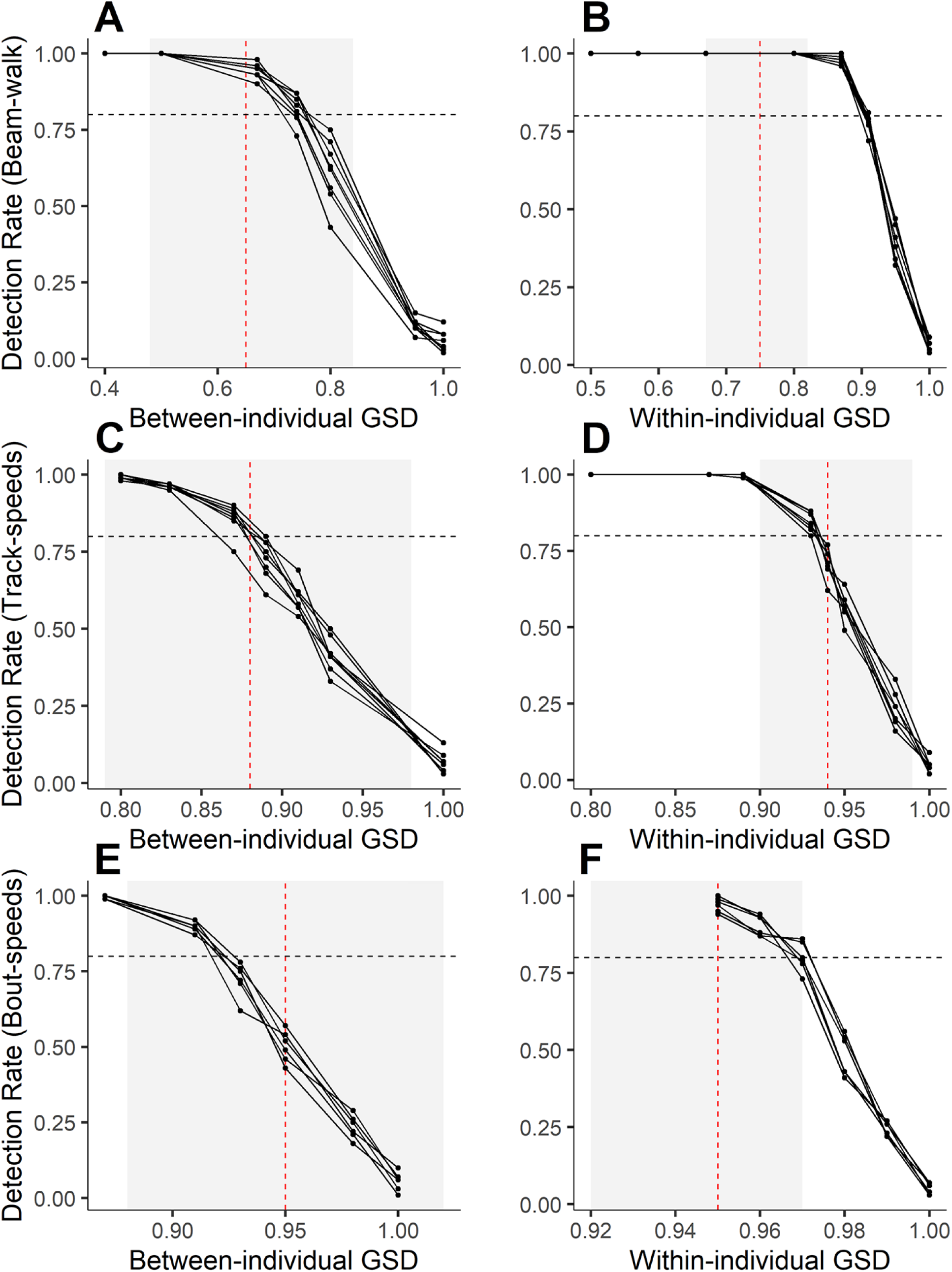
Line plots showing estimated power calculations. Panels A and B are for the beam-walk model, C and D for the track-speeds model, E and F for the bout-speeds model. X-axis shows fold difference between the two simulated phases, where decreasing values indicate a bigger difference. Y-axis shows proportion of datasets out of 100 in which a difference could be detected. Points represent values at which datasets were simulated, and lines connect points with the same value for the other variance component. The horizontal dashed line represents the 0.8 power level. The vertical red line represents the effect seen in the actual data, and the shaded interval around this represents the 95% HPDI of those model estimates.

## References

[1] Bell, A. M., Hankison, S. J., and Laskowski, K. L. The repeatability of behaviour: a meta-analysis. Animal Behaviour 77(4), 771–783 (2009).

[2] Webster, M. M. and Ward, A. J. Personality and social context. Biological Reviews 86(4), 759–773 (2010).

[3] Landeau, L. and Terborgh, J. Oddity and the ‘confusion effect’ in predation. Animal Behaviour 34(5), 1372–1380 (1986).

[4] Humphries, D. A. and Driver, P. M. Protean defence by prey animals. Oecologia 5, 285–302 (1970).

[5] Stamps, J. A. Individual differences in behavioural plasticities. Biological Reviews 91(2), 534–567 (2016).

[6] Herbert-Read, J. E., Krause, S., Morrell, L. J., Schaerf, T. M., Krause, J., and Ward, A. J. W. The role of individuality in collective group movement. Proceedings of the Royal Society B: Biological Sciences 280(1752), 20122564 (2013).

[7] Schaerf, T. M., Dillingham, P. W., and Ward, A. J. W. The effects of external cues on individual and collective behavior of shoaling fish. Science Advances 3(6), e1603201 (2017).

[8] Briffa, M. Plastic proteans: reduced predictability in the face of predation risk in hermit crabs. Biology Letters 9(5), 20130592 (2013).

[9] Houslay, T. M., Vierbuchen, M., Grimmer, A. J., Young, A. J., and Wilson, A. J. Testing the stability of behavioural coping style across stress contexts in the Trinidadian guppy. Functional Ecology 32(2), 424–438 (2017).

[10] Ghalambor, C. K., McKay, J. K., Carroll, S. P., and Reznick, D. N. Adaptive versus non-adaptive phenotypic plasticity and the potential for contemporary adaptation in new environments. Functional Ecology 21(3), 394–407 (2007).

[11] Biro, P. A. and Adriaenssens, B. Predictability as a personality trait: Consistent differences in intraindividual behavioral variation. The American Naturalist 182(5), 621–629 (2013).

[12] Urszán, T. J., Garamszegi, L. Z., Nagy, G., Hettyey, A., Török, J., and Herczeg, G. Experience during development triggers between-individual variation in behavioural plasticity. Journal of Animal Ecology 87(5), 1264–1273 (2018).

[13] Niemelä, P. T., Niehoff, P. P., Gasparini, C., Dingemanse, N. J., and Tuni, C. Crickets become behaviourally more stable when raised under higher temperatures. Behavioral Ecology and Sociobiology 73(6) (2019).

[14] Westneat, D. F., Wright, J., and Dingemanse, N. J. The biology hidden inside residual within-individual phenotypic variation. Biological Reviews 90(3), 729–743 (2014).

[15] Prentice, P. M., Houslay, T. M., Martin, J. G. A., and Wilson, J. Genetic variance for behavioural ‘predictability’ of stress response. Journal of Evolutionary Biology (2020).

[16] Cullen, D. A., Cease, A. J., Latchininsky, A. V., Ayali, A., Berry, K., Buhl, J., De Keyser, R., Foquet, B., Hadrich, J. C., Matheson, T., Ott, S. R., Poot-Pech, M. A., Robinson, B. E., Smith, J. M., Song, H., Sword, G. A., Vanden Broeck, J., Verdonck, R., Verlinden, H., and Rogers, S. M. From molecules to management: Mechanisms and consequences of locust phase polyphenism. Advances in Insect Physiology 53, 167–285 (2017).

[17] Pener, M. P. and Simpson, S. J. Locust phase polyphenism: An update. Advances in Insect Physiology 36, 1–272 (2009).

[18] West-Eberhard, M. J. Developmental plasticity and the origin of species differences. Proceedings of the National Academy of Sciences 102 (Supplement 1), 6543–6549 (2005).

[19] Roessingh, P., Simpson, S. J., and James, S. Analysis of phase-related changes in behaviour of desert locust nymphs. Proceedings of the Royal Society of London. Series B: Biological Sciences 252(1333), 43–49 (1993).

[20] Simões, P., Ott, S. R., and Niven, J. E. Associative olfactory learning in the desert locust, *Schistocerca gregaria*. Journal of Experimental Biology 214(15), 2495–2503 (2011).

[21] Simpson, S. J., McCaffery, A. R., and Hägele, B. F. A behavioural analysis of phase change in the desert locust. Biological Reviews of the Cambridge Philosophical Society 74(4), 461–480 (1999).

[22] Lochmatter, T., Roduit, P., Cianci, C., Correll, N., Jacot, J., and Martinoli, A. SwisTrack - a flexible open source tracking software for multiagent systems. In 2008 IEEE/RSJ International Conference on Intelligent Robots and Systems. IEEE, (2008).

[23] R Core Team. R: A Language and Environment for Statistical Computing. R Foundation for Statistical Computing, Vienna, Austria, (2019).

[24] RStudio Team. RStudio: Integrated Development Environment for R. RStudio, Inc., Boston, MA, (2018).

[25] Bürkner, P.-C. Advanced Bayesian multilevel modeling with the R package brms. The R Journal 10(1), 395–411 (2018).

[26] Carpenter, B., Gelman, A., Hoffman, M. D., Lee, D., Goodrich, B., Betancourt, M., Brubaker, M., Guo, J., Li, P., and Riddell, A. Stan: A probabilistic programming language. Journal of Statistical Software 76(1) (2017).

[27] Gelman, A. and Rubin, D. B. Inference from iterative simulation using multiple sequences. Statistical Science 7(4), 457–472 (1992).

[28] Pigliucci, M. and Murren, C. J. Perspective: Genetic assimulation and a possible evolutionary paradox: Can macroevolution sometimes be so fast as to pass us by? Evolution 57(7), 1455 (2003).

[29] Buhl, J., Sumpter, D. J., Couzin, I. D., Hale, J. J., Despland, E., Miller, E. R., and Simpson, S. J. From disorder to order in marching locusts. Science 312(5778), 1402–1406 (2006).

[30] Martin, J. G. A., Pirotta, E., Petelle, M. B., and Blumstein, D. T. Genetic basis of between-individual and within-individual variance of docility. Journal of Evolutionary Biology 30(4), 796–805 (2017).

[31] Rogers, S. M., Cullen, D. A., Anstey, M. L., Burrows, M., Despland, E., Dodgson, T., Matheson, T., Ott, S. R., Stettin, K., Sword, G. A., and Simpson, S. J. Rapid behavioural gregarization in the desert locust, *Schistocerca gregaria* entails synchronous changes in both activity and attraction to conspecifics. Journal of Insect Physiology 65, 9–26 (2014).

[32] Ott, S. R., Verlinden, H., Rogers, S. M., Brighton, C. H., Quah, P. S., Vleugels, R. K., Verdonck, R., and Broeck, J. V. Critical role for protein kinase a in the acquisition of gregarious behavior in the desert locust. Proceedings of the National Academy of Sciences 109(7), E381–E387 (2011).

[33] Vasquez, R. A. The influence of habitat on travel speed, intermittent locomotion, and vigilance in a diurnal rodent. Behavioral Ecology 13(2), 182–187 (2002).

[34] Bazazi, S., Bartumeus, F., Hale, J. J., and Couzin, I. D. Intermittent motion in desert locusts: Behavioural complexity in simple environments. PLoS Computational Biology 8(5), e1002498 (2012).

[35] Knebel, D., Ayali, A., Guershon, M., and Ariel, G. Intraversus intergroup variance in collective behavior. Science Advances 5(1), eaav0695 (2019).

[36] Bergmüller, R. and Taborsky, M. Animal personality due to social niche specialisation. Trends in Ecology & Evolution 25(9), 504–511 (2010).

[37] von Merten, S., Zwolak, R., and Rychlik, L. Social personality: a more social shrew species exhibits stronger differences in personality types. Animal Behaviour 127, 125–134 (2017).

[38] Jäger, H. Y., Han, C. S., and Dingemanse, N. J. Social experiences shape behavioral individuality and within-individual stability. Behavioral Ecology 30(4), 1012–1019 (2019).

[39] DiRienzo, N., Johnson, J. C., and Dornhaus, A. Juvenile social experience generates differences in behavioral variation but not averages. Behavioral Ecology 30(2), 455–464 (2018).

[40] McCune, K., Jablonski, P., im Lee, S., and Ha, R. Evidence for personality conformity, not social niche specialization in social jays. Behavioral Ecology 29(4), 910–917 (2018).

[41] Chang, C., Teo, H. Y., Norma-Rashid, Y., and Li, D. Predator personality and prey behavioural predictability jointly determine foraging performance. Scientific Reports 7 (1) (2017).

[42] Jones, K. A., Jackson, A. L., and Ruxton, G. D. Prey jitters: protean behaviour in grouped prey. Behavioral Ecology 22(4), 831–836 (2011).

[43] Richardson, G., Dickinson, P., Burman, O. H. P., and Pike, T. W. Unpredictable movement as an antipredator strategy. Proceedings of the Royal Society B: Biological Sciences 285(1885), 20181112 (2018).

[44] Limpert, E., Stahel, W. A., and Abbt, M. Log-normal distributions across the sciences: Keys and clues. BioScience 51(5), 341 (2001).

[45] Tatliyer, A., Cervantes, I., Formoso-Rafferty, N., and Gutiérrez, J. P. The statistical scale effect as a source of positive genetic correlation between mean and variability: A simulation study. G3; Genes|Genomes||Genetics 9(9), 3001–3008 (2019).

